# Canonical features of human antibodies recognizing the influenza hemagglutinin trimer interface

**DOI:** 10.1101/2020.12.31.424868

**Authors:** Seth J. Zost, Jinhui Dong, Iuliia Gilchuk, Pavlo Gilchuk, Natalie J. Thornburg, Sandhya Bangaru, Nurgun Kose, Jessica A. Finn, Robin Bombardi, Cinque Soto, Rachel Nargi, Ryan P. Irving, Naveenchandra Suryadevara, Jonna B. Westover, Robert H. Carnahan, Hannah L. Turner, Sheng Li, Andrew B. Ward, James E. Crowe

## Abstract

Broadly reactive antibodies targeting the influenza A hemagglutinin (HA) head domain are thought to be rare and to require extensive somatic mutations or unusual structural features to achieve breadth against divergent HA subtypes. Here we describe common genetic and structural features of diverse human antibodies from several individuals recognizing the trimer interface (TI) of the influenza HA head, a recently identified site of vulnerability^1–3^. We examined the sequence of TI-reactive antibodies, determined crystal structures for TI antibody-antigen complexes, and analyzed the contact residues of the antibodies on HA to discover common genetic and structural features of TI antibodies. Our data reveal that many TI antibodies are encoded by a light chain variable gene segment incorporating a shared somatic mutation. In addition, these antibodies have a shared acidic residue in the heavy chain despite originating from diverse heavy chain variable gene segments. These studies show that the TI region of influenza HA is a major antigenic site with conserved structural features that are recognized by a common human B cell public clonotype. The canonical nature of this antibody-antigen interaction suggests that the TI epitope might serve as an important new target for structure-based vaccine design.

## INTRODUCTION

Influenza A virus (IAV) is one of the most common causes of severe lower respiratory illness in humans and exhibits a wide antigenic diversity in circulating field strains. Seasonal epidemics with H1 and H3 IAV subtypes occurs yearly and other zoonotic IAVs with H1, H3, H5, H6, H7, H9 and H10 HAs cause outbreaks of human infections sporadically^4,5^. Incomplete matches of the seasonal IAV vaccine strains with the antigenically drifted viruses causing epidemics lead to vaccine ineffectiveness and contribute to severe influenza seasons ^6–8^.

A high priority for current IAV vaccine development is the development of vaccine antigens that induce broad and protective immune responses^9,10^. Antibodies to the hemagglutinin (HA) stem domain can exhibit heterosubtypic recognition patterns^11–24^. However, the accessibility of the stem domain may be reduced on the virion particle, and stem-specific antibodies exhibit mostly a moderate protective capacity. The HA head domain is immunodominant and is the target of most antibody responses induced by IAV vaccine or infection^25–29^, but head domain specific antibodies often recognize a narrow spectrum of IAVs. Recently, several groups have identified an entirely new class of human antibodies recognizing a highly conserved region on the HA head domain, the TI region^1–3^. The few monoclonal antibodies (mAbs) reported have broad recognition patterns for diverse IAVs, are non-neutralizing, and protect in animal models^1–3^. The critical HA residues recognized by the TI mAbs reported to date remain conserved across most subtypes of influenza A viruses, suggesting that the TI site might serve as an attractive new antigenic target for epitope-based universal IAV vaccine design. However, designing vaccine antigens based on recognition of unusual antibodies isolated only from rare individuals is not desirable, since universal influenza vaccine antigens should possess the capacity to induce broadly protective antibodies in a large number of individuals with diverse exposure histories. Here, we sought to isolate more TI mAbs and determine if TI antibodies possess any common features that are present in human repertoires. Remarkably, we found that many TI antibodies are members of a common public B cell clonotype with canonical genetic and structural features, suggesting that most human subjects have the capacity to make TI antibodies with a minimal number of somatic mutations needed to accomplish near-universal influenza A recognition.

## RESULTS

### Identification of broadly reactive human TI mAbs in a panel of H5 HA-specific mAbs

We previously reported isolation of H5 HA-specific human antibodies from otherwise healthy subjects who had received an A/Vietnam/1203/2004 H5N1 (VN/04) subunit vaccine^30^. Here we examined the reactivity of some of the H5-reactive mAbs to determine if any exhibited heterosubtypic breadth of recognition for diverse HA subtypes. To investigate the breadth, we tested purified IgGs for each for binding activity to HA from different IAV subtypes; all HA proteins used were recombinant trimers. MAbs designated H5.28 and H5.31 exhibited breadth of binding to recombinant HAs belonging to group 1 (H1, H2, H5, H6, H8, H9, H11 and H12) and group 2 (H3, H4, H7, H10, H14 and H15) viruses (**Fig. 1a**). DNA copies of the wild-type H5.28 and H5.31 variable regions were synthesized and recombinant forms of IgG proteins were expressed; hybridoma-generated antibody (designated H5.28 or H5.31) was used for the assays unless the recombinant form is specified (designated as rH5.28 or rH5.31). As expected, the rH5.28 and rH5.31 IgGs showed a similar binding pattern to the corresponding hybridoma-produced IgG proteins. Neither H5.28 nor H5.31 had hemagglutinin-inhibiting or neutralizing activity for a reverse genetics-derived VN/04 H5N1 virus when tested in concentrations as high as 10 μg/mL as the original hybridoma derived IgG nor as recombinant purified IgG molecules made in 293F or Chinese hamster ovary (CHO) cells (**Extended Data Fig. 1**). Despite the fact that these antibodies did not neutralize virus *in vitro*, we tested whether these mAbs protected against weight loss and death in mice following a stringent lethal challenge with a mouse-adapted A/California/04/2009 (H1N1pdm) strain. BALB/c mice (n=10 per group) were administered 1, 3 or 10 mg/kg of H5.28 or H5.31 IgG or a similarly prepared control antibody by the intraperitoneal route, and then challenged by the intranasal route 24 hours later with a lethal dose of virus. H5.28 or H5.31 mediated dose-dependent protection against mortality and protection against severe weight loss when administered prophylactically at three tested doses. Protection with low dose mAb treatment was comparable to that of high-dose oseltamivir given at a high daily dose of 30 mg/kg/day on days 1 to 5 after virus inoculation. Mice treated with H5.28 (**Fig 1b**), or H5.31 (**Fig 1c**) (n=10 for each group) showed protection from weight loss after virus challenge in a dose-dependent manner whereas mice treated with PBS or the 2D22 control antibody succumbed to infection. These results indicate the ability of mAb H5.28 or H5.31 to protect *in vivo* against lethal virus challenge against a virulent influenza A virus strain.

**Fig. 1.**
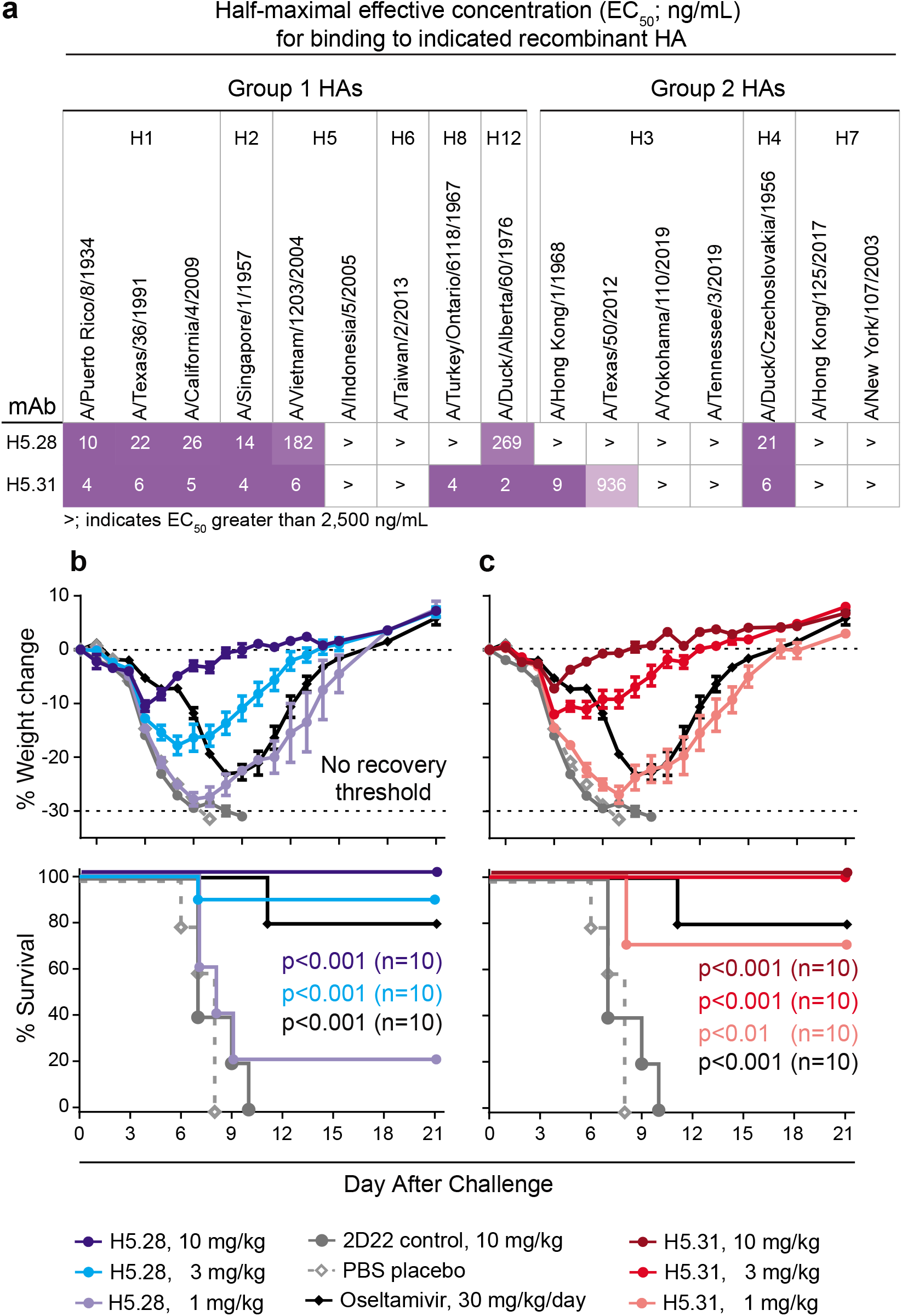
MAb H5.28 and H5.31 cross-react broadly and protect *in vivo*. **a,** ELISA to determine strength of binding of H5.28 or H5.31 to a panel of recombinant HAs. EC50 values for binding (ng/mL) are shown in each square. The purple-white color scale indicates strength of binding, with darker squares indicating strong binding. **b-c**, Body-weight changes and survival in mice that received prophylactic treatment with mAb H5.28 (**b,** blue lines) or mAb H5.31 (**c,** red lines) at three different doses. Mice were challenged intranasally with a lethal dose (2,200 CCID50) of A/California/04/2009 virus 24 hrs after prophylactic administration. A negative control group was treated with the anti-dengue virus mAb DENV 2D22 (solid grey lines), while a positive control group was treated with oseltamivir daily for 5 days (black lines). An additional control group received a PBS placebo (dashed grey lines). For weight loss curves, error bars show the standard error of the mean (SEM). The Mantel-Cox log-rank test was used to compare each survival curve to the negative control DENV 2D22 treatment group, and associated p-values from a comparison to the control treatment group are shown.

To identify the specific epitope recognized by these mAbs, we performed hydrogen deuterium exchange mass spectrometry (HDX-MS) experiments. We used a monomeric head domain of H5 (based on strain VN/04) to identify peptides on the surface of HA that are occluded following binding of H5.28. We found that H5.28 Fab reduced deuterium labeling of peptides comprising residues 96-105, 136-147, and 217-233 (H3 structure numbering, **Fig 2a and Extended Data Fig. 2**). From the HDX-MS studies, we anticipated that H5.28 or H5.31 binding to the HA trimer destabilizes the trimeric interface of native HA. To directly examine the effect of these Fabs on the HA trimer, we performed negative-stain electron microscopy (nsEM) of HA (uncleaved H1 A/California/04/2009 stabilized with a GCN4 trimerization motif [H1 HA0]) in complex with either H5.28 or H5.31 Fab incubated for different lengths of time. Native H1 HA0 trimer remained in its trimeric conformation during nsEM sample preparation (**Fig 2b**). In contrast, we observed that upon exposure to H5.28 or H5.31 even for 20 seconds (the shortest time point that could be tested), the HA0 trimers quickly degraded into Fab-bound monomeric HA, with only a small fraction of Fab-free HA remaining in a trimeric conformation (**Fig 2b**). Despite extensive trials, the intermediate stage of this structural change could not be obtained, apparently due to the rapid transformation of the HA0 from trimeric to monomeric states induced by antibody binding. These results demonstrate that H5.28 and H5.31 bind the uncleaved HA0 trimer, and then dissociate the trimer *in vitro* (**Fig 2b**). The ability to selectively disrupt HA0 trimers on the surface of infected cells and consequently inhibit cell-cell spread is consistent with this *in vitro* phenomenon. Both mAbs H5.28 and H5.31 bound preferentially to uncleaved HA (with reduced binding to cleaved HA) on the surface of HA-transfected cells, while a recombinant form of a representative stem domain antibody bound to both forms well (**Fig 2c, Extended Data Fig. 3**).

**Fig. 2.**
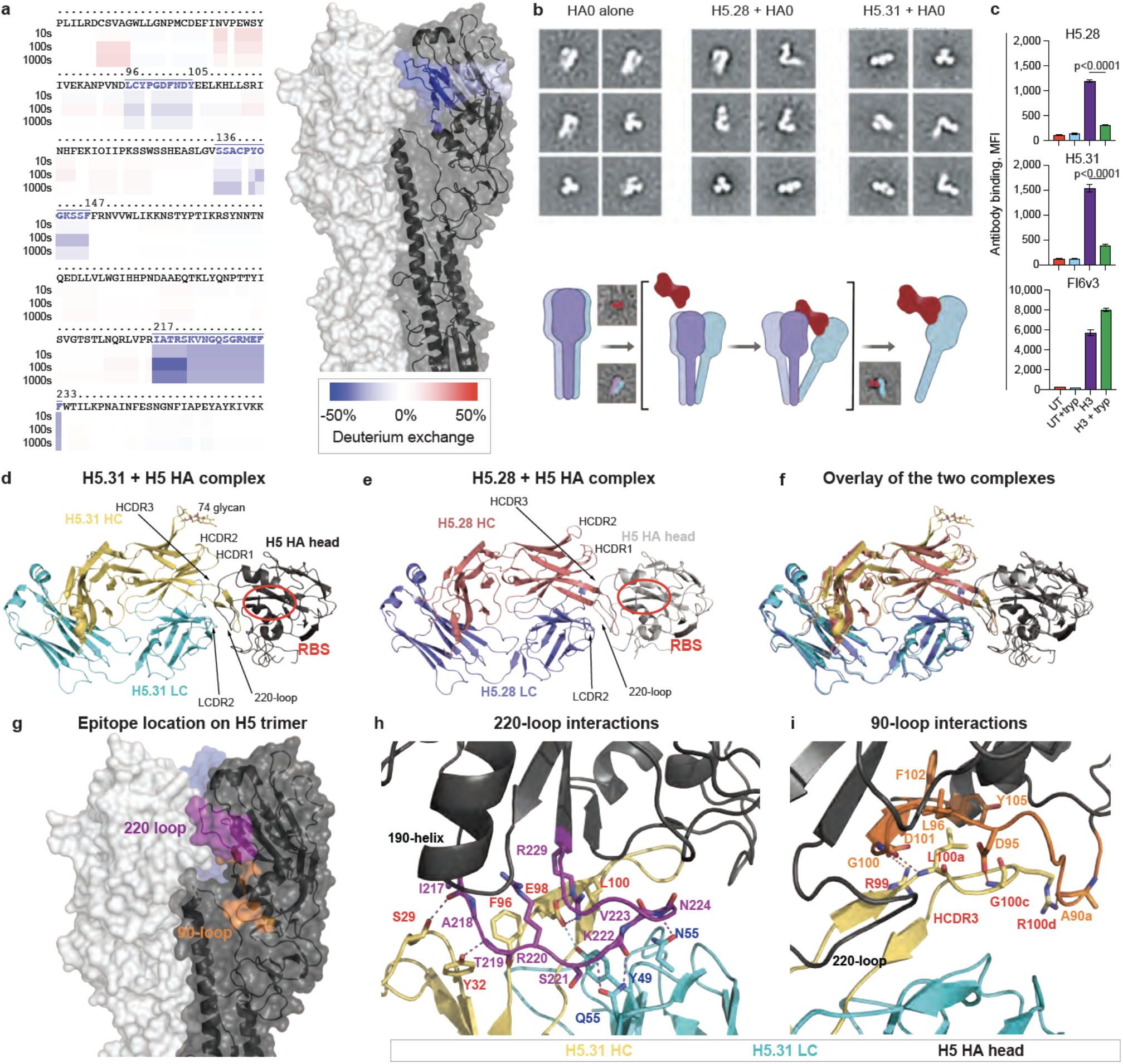
Structural and functional characterization of mAb H5.28 and H5.31 binding to HA. **a**, HDX-MS with H5.28 Fab and H5 HA head domain identified a putative epitope for H5.28 at the trimer-interface. The amino acid sequence of the H5 head domain is shown with a ribbon diagram indicating differences in deuterium uptake. Blue colors indicating slower deuterium exchange in the presence of H5.28 Fab, while red colors indicate faster deuterium exchange in the presence of H5.28 Fab. Data are shown for 10s, 100s, and 1000s of deuterium labeling. The three major peptides are colored blue on an H5 trimer. **b**, Selected 2D class averages of H1 HA trimer (A/California/04/2009) alone or after a 20-second incubation with H5.28 (middle) or H5.31 Fab (right). All of the Fabs complexed with HA were in monomeric form, while a few apo HA trimers were observed. Below is a cartoon illustration showing that H5.28 or H5.31 Fab (magenta) results in dissociation of native HA trimer (grey/blue/lavender), as visualized in the negative-stain EM data. **c**, H5.28 and H5.31 preferentially bound to uncleaved HA expressed on the cell surface. In contrast, a recombinant form of the stem mAb FI6v3 preferentially bound to cleaved HA. p-values for the comparison between binding to cleaved and uncleaved HA were computed using an unpaired two-way Student’s t-test. **d**, Crystal structure of H5.31 in complex with VN/04 HA. H5.31 heavy chain is colored in gold, the light chain in cyan, and H5 head domain in dark grey. A glycosylation site on H5.31, N74 on the H5.31 DE-loop, is labeled and shown as sticks, and two fitted NAG residues of the glycan are shown also as sticks. The 220-loop of the VN/04 HA head domain is labeled, and the receptor binding site is highlighted with a red circle. **e**, Crystal structure of H5.28 in complex with VN/04 HA. H5.28 heavy chain is colored in salmon, the light chain in lavender, and H5 HA head domain in light grey. **f,** Superimposition of the two complex structures. Structural variations can be seen at HCDR1s, HCDR2s, and heavy chain DE-loops. **g**, The HA epitope recognized by H5.31 is mapped onto the surface of one protomer of the VN/04 HA trimer. The mapped protomer is colored dark grey, the 220-loop in purple, and the 90-loop in orange. The other two protomers are colored in light grey and blue respectively. **h-i**, Structural details of the H5.31/VN/04-HA complex. **h**, Interactions of H5.31 with 220-loop of the HA head domain, with the HCDR3 E98/LCDR2 Y49 interaction shown with blue dashes. Relevant residues of the HA head domain are labeled in purple, those of the heavy chain in red, and those of the light chain in blue. **i**, Interactions of H5.31 HCDR3 with 90-loop and its C-terminal β-strand. Relevant residues of the HA head domain are labeled in orange, those of the heavy chain in red, and those of the light chain in blue.

To determine the molecular details of the interaction of H5.28 and H5.31 with the TI site, crystal structures of the H5.31 or H5.28 Fabs and their complexes with the HA head domain from VN/04 were determined at 3.00 Å or 4.00 Å resolution, respectively (**Table S1**). The complex structures revealed that H5.28 and H5.31 recognize an epitope in the TI region very similar to that of the previously reported FluA-20 and S5V2-29 mAbs (**Fig 2d-f**). Antibodies H5.31 and H5.28 are clonally related siblings from one human subject with identical HCDR3 sequences and only several amino acid variations in their light and heavy chains. The overlay of the two crystal structures showed that H5.28 binds to the HA head domain in the same general manner as H5.31, although some regions of H5.31, such as the heavy chain DE loop, HCDR1, and HCDR2 deviate from that of H5.28 (**Fig 2f**). The superposition of the variable domain and HA head domain of H5.31-HA onto those of H5.28-HA results in a Cα RMSD of only 0.71 Å. The epitope recognized by H5.31/H5.28 on VN/04 maps onto one HA1 protomer of the H5N1 HA trimer (PDB ID: 4BGW) (**Fig 2g).** This epitope can be divided into two regions: the 220–loop of the receptor-binding domain (residues 217 - 224, and residue 229, H3 structure numbering) and a second region at the 90–loop (**Fig 2g**). The sequences of the 220–loop of influenza A HA are relatively conserved, thus recognition of this region by H5.31/H5.28 partly explains the binding breadth of the two mAbs. In addition, the epitope recognized by H5.31 and H5.28 is inaccessible for mAbs to bind in the closed HA trimeric form (**Fig 2g**). If we superimpose the head domain of H5.31/H5-HA complex onto H5N1 HA trimer (PDB ID: 4BGW), the H5.31 heavy chain variable domain would occupy the space of the head domain of another adjacent HA protomer, *e.g*., in the closed trimer the head domain of an adjacent HA1 protomer clashes with the variable domain of H5.31 when bound. Therefore, the HA trimer must make large structural rearrangements from its classic static conformation to expose the TI epitope in order for H5.31 to bind the trimer. This finding suggests the HA trimer has large conformational fluctuations in its quaternary structure, at least for HA1 including the head domains, even at neutral pH.

In the H5.31/H5-HA structure, mAb H5.31 interacts with the HA 220-loop using HCDR3 and LDCR2 residues (**Fig 2h**). There are 6 hydrogen bonds (H-bond) or salt bridges between the 220–loop and the HCDRs. The highly conserved HA 220–loop residue R229 forms a salt bridge with the mAb H5.31 HCDR3 residue E98, and the salt bridge is mostly buried, emphasizing its importance in contributing to the binding free energy. Notably, LDCR2 residue Y49 forms an H-bond with the sidechain of HCDR3 residue E98, assisting E98 to be well-positioned to interact with the 220–loop residue R229. All of those H-bonds are formed between the 220–loop mainchain oxygen or nitrogen atoms and side chains of the mAb, and consequently this mode of H-bond formation may contribute to the great breadth of the mAb. A hydrophobic interaction between mAb H5.31 residue L100 and the 220–loop residue V223, and cation-π interaction between 220–loop R220 and HCDR3 F96, also may contribute to the tight binding of H5.31. In addition, the tip of the elongated HCDR3 makes more contact with the 90–loop epitope, in which residue L100a (Kabat numbering) seems to play the major role (**Fig 2i**). The L100a sidechain is surrounded by a hydrophobic pocket formed by HA residues L96, F102, and Y105, and its mainchain nitrogen forms a H-bond with the HA G100 mainchain oxygen. Peripheral to these L100a-Ag interactions, there are several polar interactions, such as a polar interaction between the HA D95 residue sidechain and the HCDR3 G100c mainchain nitrogen and salt bridge between D101 (HA) and R99 (HCDR3). In summary, H5.31 recognizes the HA head domain mainly by interacting with the HA 220–loop, including one salt bridge with a conserved arginine residue and 5 H-bonds with mainchain atoms of the 220–loop, and the tip of the elongated HCDR3 provides additional interaction with the 90–loop and its C-terminal β strand.

Comparison of the crystal structures of 4 mAb complexes with the TI site revealed conserved features. First, a negatively charged residue (D or E) is always present in the HCDR3, forming strong salt bridges with the highly conserved arginine residues in the 220–loop (R229 in H3 structure numbering), and residue Y49 from the LDCR2s positions the negatively-charged residue via a H-bond for the formation of the salt bridge. Second, two residues from LDCR2 (N53 and Q55) form H-bonds with mainchain atoms of 220–loop. Lastly, the tips of the HCDR3s make additional contacts with the 90–loop and adjacent structural elements.

Interestingly, heavy chain DE loop in the framework region 3 (FR3) of H5.31 has a potential glycosylation site at residue N74 (Kabat numbering, sequence motif: N^74^ASN^77^), and two NAG residues can be fit into electron density around residue N74. The apparent molecular weight of H5.31 (but not H5.28) in SDS-PAGE gels shifted to a lower value with PNGase or Endo H enzymatic digestion, but the binding pattern of the glycosylated and de-glycosylated forms of H5.31 could not be distinguished in binding to HA (data not shown). Therefore, H5.31 is glycosylated in FR3, but without apparent functional alternation due to this modification.

Since the H5.28, H5.31, Flu-A20, and S5V2-29 mAbs are encoded by light chains with common features, we tested the hypothesis that a sequence signature associated with use of this gene could be used to identify new TI-specific antibodies in the antibody variable gene repertoires of a subject during acute natural infection. We studied the response of an otherwise healthy subject with exposure to diverse influenza vaccines who presented with acute laboratory-confirmed H3N2 virus respiratory infection in August 2017. For comparative purposes, we used deep sequencing to profile the B cell repertoire of this individual at various time points before or after natural infection. Sequencing timepoints included both healthy state baselines as well as responses to influenza vaccination (**Fig 3a**). At a timepoint approximately one week into the natural H3N2 infection, we obtained PBMCs, isolated plasmablasts (**Extended Data Fig. 4**), and performed single-cell sequencing of expressed paired heavy and light chain mRNA (sc-V_H_:V_L_Seq) on ~20,000 plasmablasts. We synthesized cDNA from a subset of recovered pairs of antibody genes and expressed the heavy-light chain pairs individually in small-scale CHO cell culture and then purified IgG from cell supernatants with Protein G affinity resin. Purified recombinant antibodies were tested by ELISA for binding to diverse HAs (Group 1: H1, H5; Group 2: H3, H7), and by neutralization of a representative H3N2 wild-type virus corresponding to a recent H3N2 vaccine strain (A/Texas/50/2012). 16 of the antibodies exhibited heterosubtypic reactivity (binding to more than one HA subtype) and HA protein specificity.

**Fig. 3.**
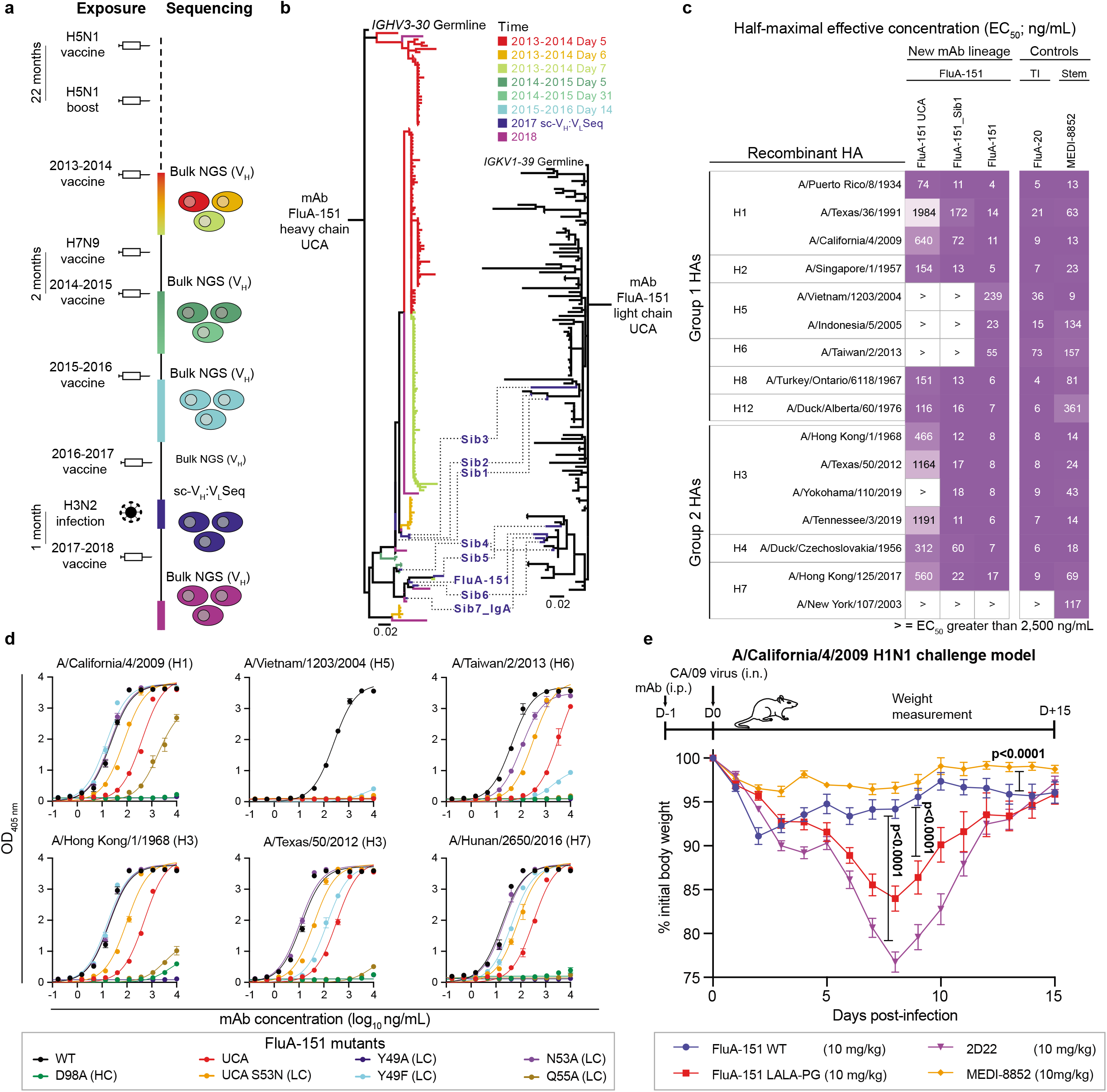
Identification and functional characterization of a new TI mAb lineage recalled in the response to vaccination and natural infection. **a**, Timeline showing the vaccination history of the research subject, with exposures and repertoire sequencing indicated. **b**, Phylogenetic trees showing the FluA-151 lineage over four years of vaccination and infection, with branch colors corresponding to sequencing timepoint. At left, the heavy chain phylogeny for FluA-151 is color-coded based on year of vaccination and days post-vaccination. At right, the light chain phylogeny for FluA-151 is shown. Paired heavy-light sequences identified by single-cell sequencing are shown with dashes connecting the heavy chain and light chain trees. **c**, Binding of TI mAb FluA-151, a clonally-related mAb (FluA-151_Sib1), and the inferred unmutated common ancestor of FluA-151 (FluA-151 UCA) to a panel of recombinant HAs. The previously described TI mAb FluA-20 and a recombinant version of the broadly-reactive stem mAb MEDI-8852 are shown for comparison. FluA-20 and FluA-151 did not bind measurably to the A/New York/107/2003 HA, which has a 220-loop deletion. **d**, ELISA binding of FluA-151 point mutants to HAs from different strains. Points and error bars represent the mean ± SD of technical triplicates. Experiments were repeated twice, with data from one representative experiment shown. **e**, FluA-151 protection from weight loss in a sublethal H1N1 challenge model. Mice were passively transferred with either FluA-151 WT (blue), FluA-151 LALA-PG (red), the positive control anti-stem mAb MEDI-8852 (orange), or the negative control anti-dengue mAb 2D22 (purple) one day prior to intranasal challenge with a sublethal dose of A/California/04/2009. For weight loss curves, error bars show the standard error of the mean (SEM). Statistical comparisons between treatment groups were performed using a repeated measurements two-way analysis of variance (ANOVA) with Tukey’s correction for multiple comparisons.

These heterosubtypic antibodies included siblings of FluA-20. However, we also identified other heterosubtypic mAbs that used the *IGKV1-39* light chain characteristic of FluA-20, H5.28, and H5.31. Sequence alignment for one of these new mAbs, designated FluA-151, showed that 7 additional members of the clonal lineage for that antibody (which we designated Sibs 1-7) were also present in the collection of over 4,000 plasmablast paired heavy-light chain sequences (indicated as Sibs in **Fig 3b**). We searched for FluA-151-like sequences in the collection of *all* antibody repertoire sequences for that donor obtained over the four-year period 2014-18 and found 178 additional somatic variants of the heavy chain and 99 additional variants of the light chain (**Fig 3b**). We constructed lineages of the clonotype, showing all corresponding heavy and light chain sequences, indicating the year and day after vaccination for the sample from which the variant was obtained (**Fig 3b**). The lineage included a diverse range of sequences and members of this lineage appeared over multiple years of responses to vaccination. We expressed FluA-151, its inferred UCA (FluA-151 UCA), and the Sib 1 variant (FluA-151_Sib1) and tested the heterosubtypic breadth of these related antibodies. The UCA had a relatively broad pattern of binding, recognizing HAs from Group 1 (H1, H2, H8, H12), and Group 2 (H3, H4, and H7) (**Fig 3c**). The intermediate FluA-151 Sib1 acquired recognition of a 2019 H3N2 strain and improved the EC50 value of binding for most Group 2 strains. The fully mature FluA-151 mAb was even broader, acquiring binding capacity for H5 and H6. These data show that the founder clone of the lineage was influenza HA-reactive and had substantial heterosubtypic breadth, and somatic mutations that occurred during elaboration of the lineage further enhanced heterosubtypic breadth. Notably, both FluA-151 and FluA-20 did not exhibit detectable binding to the HA from A/New York/107/2003, which has a deletion in the 220 loop, indicating that FluA-151 was likely a TI mAb.

We used our knowledge of the contact residues of other TI mAbs to define FluA-151 residues critical for epitope recognition using mutagenesis^2,3^. We previously demonstrated that alanine substitutions at D98 of the FluA-20 heavy chain or Y49 and Q55 of the FluA-20 light chain abrogated binding of FluA-20. Introducing these same mutations into FluA-151 greatly reduced binding against all HAs tested (**Fig. 3d**). FluA-20, FluA-151, S5V2-29, H5.28 and H5.31 all share a S53N somatic mutation in the LCDR2, and introducing the S53N substitution into the FluA-151 UCA improved binding 5-10 fold against most HAs tested (**Fig 3d**), providing evidence that this substitution improves binding and explaining the convergent evolution of the S53N substitution across multiple TI mAb lineages isolated from multiple donors. In the background of the mature FluA-151 mAb, the N53A substitution had a modest impact on binding with the exception of binding to the VN/04 HA (**Fig. 3d**). These data indicate that in the mature form of the mAb the importance of N53-mediated interactions ranged from non-essential to critical depending on variations at the TI epitope between different strains. In contract to the dramatic effect of the Y49A substitution, the more conservative Y49F substitution did not affect the binding to some HAs but dramatically reduced binding to H5 and H6 HAs. Taken together, these results identify that FluA-151 shares many critical contact residues with FluA-20 and demonstrates that the S53N mutation is a common solution to enhance binding of multiple TI mAbs.

We next assessed the ability of FluA-151 to protect mice from weight loss following viral challenge in a sub-lethal model using an A/California/04/2009 H1N1 virus. One day prior to infection, we passively transferred mice with either FluA-151 WT or a FluA-151 LALA-PG Fc variant in which three mutations ablate Fc effector function^31^ (**Fig. 3e**). Mice that received the negative control mAb 2D22 lost more than 20% initial body weight but subsequently recovered. In contrast, animals treated with the positive control anti-stem mAb MEDI-8852 were completely protected from weight loss. While FluA-151 WT protected mice from major weight loss, mice who received the FluA-151 LALA-PG variant lost significantly more weight (**Fig. 3e**), indicating that in the case of FluA-151, Fc-mediated effector function likely mediates protection from severe weight loss associated with influenza infection.

To further understand structural determinants of the molecular recognition of the TI site by the H5.31, FluA-20, and S5V2-29 antibodies, we overlaid the crystal structures of the three antigen-antibody complexes. The complexes were remarkably similar on a global basis (**Fig. 4a**). The critical contact residues for each of the three Fabs (each derived from independent subjects) were identical. The Fabs shared a critical contact in the HCDR3 loop with an acidic (D or E) residue at Chothia position 98 contacting the R229 residue on HA (**Fig. 4b**). The Fabs also shared three critical bonds on two residues made by the light chain, at HA residues 222 and 224 (**Fig. 4c**). In addition, for all antibodies the light chain Y49 residue formed a hydrogen bond with the D or E residue at position 98 in the heavy chain (**Fig. 4c**). This interaction may be required for optimal alignment of residues for TI epitope recognition and might explain the effect of the Y49F substitution on the ability of FluA-151 to recognize some HAs (**Fig. 3d**). We aligned the antibody variable region sequences of FluA-20^2^, FluA-151, H5.28, H5.31 and the SVV2-29, S5V2-52, S1V2-37 and S1V2-58 antibodies^3^ (**Fig. 4d**) and observed a canonical sequence pattern, with a motif comprising: 1) Use of the *IGKV1-39* light chain gene, 2) conservation of germline-encoded Y49 and Q55 residues in the light chain, 3) introduction of an S53N somatic mutation in the light chain CDR2, and 4) inclusion of an acidic (D or E) residue in the HCDR3 at Chothia position 98 (which is encoded by non-templated nucleotides in the N1 region of the V_H_-D_H_ gene junction added during recombination). Although FluA-20, S5V2-29, and H5.31 all shared these sequence features that mediate interactions with the HA 220-loop, S5V2-29 and H5.31 have longer HCDR3s and as a result make more contacts with the HA (**Extended Data Fig. 5**). Clashes introduced by these additional contacts likely explain why these mAbs have more restricted reactivity. In contrast, mAbs FluA-20 and FluA-151 have shorter HCDR3s that likely avoid contacts with more variable HA residues.

**Fig. 4.**
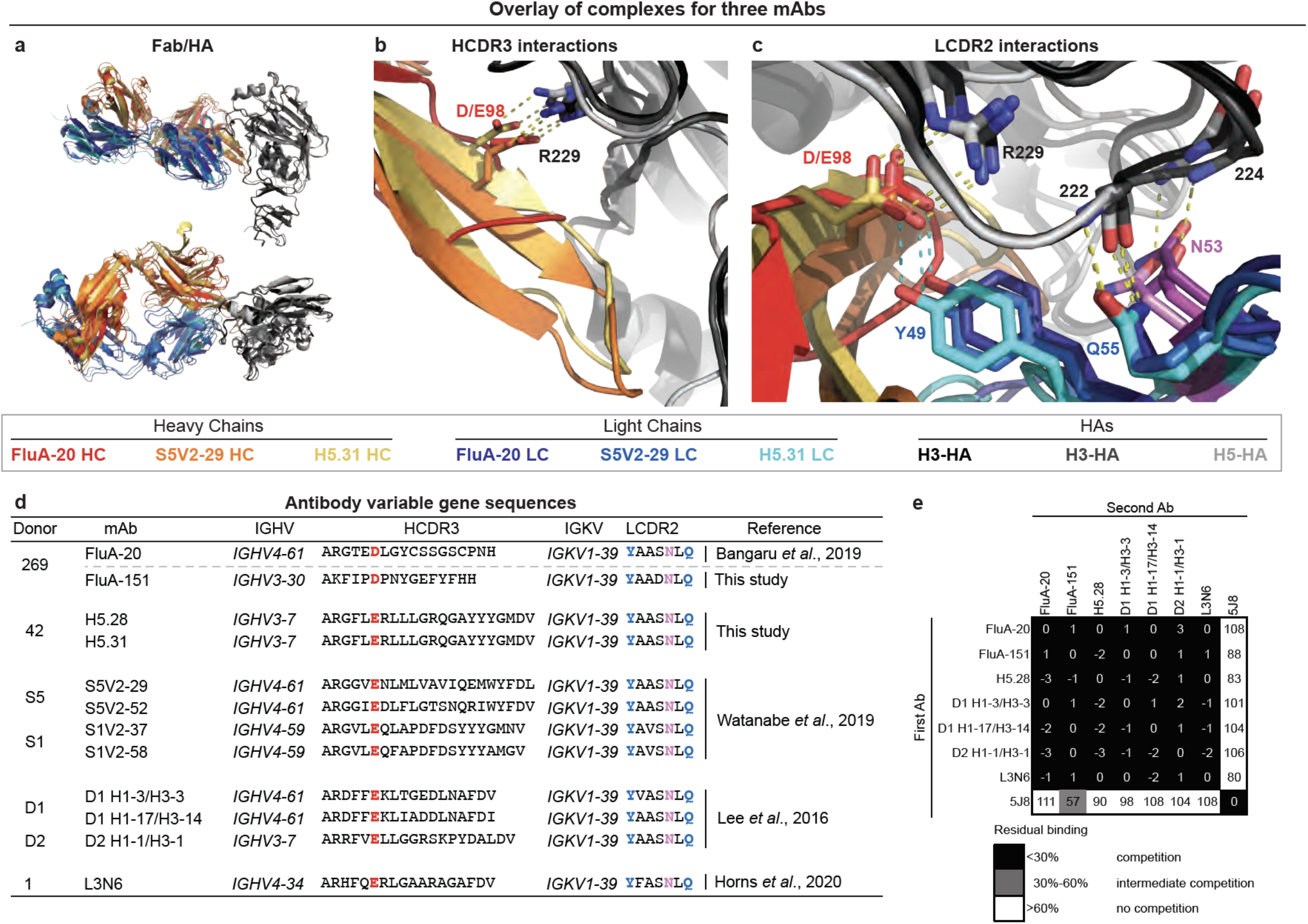
Structural and sequence alignments of TI mAbs reveal common features of epitope recognition. **a-c**, Structural alignment of the Fab:HA head domain complexes of FluA-20, S5V2-29, and H5.31, with the HA head domains aligned to one another. **a**, View of the structural alignment from the side (upper image, panel **a**) and top (lower image, panel **a**). **b**, Despite differences in HCDR3 length, FluA-20, S5V2-29, and H5.31 all contact HA R229 using a D or E at heavy chain position 98. **c**, FluA-20, S5V2-29, and H5.31 germline-encoded light chain residue Y49 makes hydrophobic contacts and a hydrogen bond with D98 or E98 of the heavy chain, while germline-encoded Q55 makes hydrogen bond contacts with the main-chain amide and carbonyl groups of HA residue 222. The shared somatic mutation S53N introduces an additional hydrogen bond with the main-chain amide of HA residue 224. **d**, Sequence alignment of previously reported and newly reported TI mAbs identifies common recognition motifs, including a shared acidic residue at position 98 in the HCDR3, a common light chain (*IGKV1-39*), germline-encoded light chain contact residues shared by all mAbs, as well as a common light chain S53N somatic mutation. The interaction between HCDR3 E98 and LCDR2 Y49 is shown with blue dashes. **e,** Biolayer interferometry-based competition data demonstrates that previously reported TI mAbs strongly compete with one another for binding to A/California/04/2009 HA trimer, but do not compete with the RBS-binding mAb 5J8.

We also considered that previous investigators had reported some human HA-specific antibodies identified by proteomic sequence analysis of serum antibodies that also displaced HA protomers during binding, indicating that these mAbs likely recognized an epitope at the interface between monomers^1^. We aligned the amino acid sequences of those three antibodies (D1 H1-3/H3-3, D1 H1-17/H3-14, and D2 H1-1/H3-1) and found that those sequences perfectly fulfilled the TI sequence motif described above. These antibodies had been assigned putatively as binding to an occluded epitope near the trimer interface of the TI HA head domain based on a low-resolution EM structure and interference with binding of a trimer-specific antibody^1^ (**Extended Data Fig. 6**). However, recombinant forms of the D1 H1-3/H3-3, D1 H1-17/H3-14, and D2 H1-1/H3-1 antibodies competed for binding to trimeric HA with TI-specific antibodies but not with the receptor binding domain specific antibody 5J8 (**Fig. 4e**). These mAbs also competed with TI mAbs for binding to a monomeric HA head domain (**Extended Data Fig 6c**). In addition, a separate group has recently described an antibody lineage with heterosubtypic breadth discovered using single B cell sequencing^32^. This antibody, L3N6, also exhibits TI mAb sequence features (**Fig. 4d**) and competes for binding with known TI mAbs (**Fig. 4e**). Together these data suggest these mAbs previously described by other groups also are in fact TI-specific antibodies.

## DISCUSSION

In this work we use fine structural characterization of naturally occurring broad human mAbs that bind the HA trimer interface to reveal canonical features of common human antibodies that bind a highly conserved site of vulnerability on influenza HA. These TI antibodies are common in human B cell repertoires, and they exhibit stereotypical features. Crystal structures of diverse TI mAbs with common light chain features reveal how the public clonotype we describe interacts with the highly conserved TI region and we demonstrate that a shared somatic mutation improves binding, providing an explanation for the convergent evolution observed across mAb lineages isolated from multiple donors. We also demonstrate that previously described mAbs with broad reactivity recognize the same TI epitope using these shared sequence features. We note that previous investigators have determined the structure of H2214, a TI mAb that contacts many of the same HA residues but does not possess the canonical features we describe here,^3^ and that a separate group has described an antibody isolated using phage display that recognizes the opposing face of the TI epitope and binds H7 and H15 HAs^33^. Thus, we conclude that in addition to the common recognition motif we identify, there may be additional modes of TI epitope recognition that have yet to be defined. When administered prophylactically or therapeutically, TI antibodies protect mice against challenge with diverse IAV strains^1–3^, suggesting these mAbs may play an important role in protecting individuals from severe disease during normal seasonal circulation of influenza A viruses. While mass spectrometry approaches suggest these mAbs are highly prevalent in the serum of some individuals^1^, quantifying the levels of these TI antibodies in humans and their contribution to protection is an important area of future work.

While these TI mAbs do not measurably neutralize virus *in vitro*, passive transfer of TI mAbs can prevent severe disease in animal models, raising important questions remain regarding how this epitope is recognized in the context of an immune response. We showed that HA cleavage reduces mAb recognition of the TI epitope (**Fig. 2c and Bangaru *et al***.^2^), but other factors also can affect TI epitope accessibility. Previous work identified mouse mAbs that bound better to HA from acid-treated viruses, suggesting they recognized pH-sensitive cryptic epitopes^34^. One of these mAbs, Y8-10C2, binds an occluded epitope at the TI based on escape mutation epitope mapping^35^. Importantly, Y8-10C2 also selected escape mutations in the HA stem region, demonstrating that mutations in this region likely affect trimer stability and epitope accessibility. We have previously described the structure of a mAb, H7.5, that partially displaces HA monomers to recognize a cryptic epitope near the TI, but binds in a different pose from the TI mAbs we describe here^36^. Finally, others have demonstrated that HA hyperglycosylation focuses mouse antibody responses to TI epitopes^37^. Taken together, these findings indicate that 1) the HA is a less static structure than is often assumed, 2) multiple factors can influence the accessibility of TI epitopes, and 3) TI epitopes are commonly recognized during antibody responses induced by either natural infection or vaccination.

The work we present here also has implications for characterization of recurring structural motifs that facilitate antibody recognition of antigens. Historically, most B cell repertoire sequencing has focused on examination of the heavy chain, especially on the HCDR3 region, which dominates most antigen-antibody interactions. In one example, the most common class of human antibodies to the IAV HA stem region (encoded by the *IGHV1-69* gene) exhibits promiscuous paring with various light chains^38^. Members of the class of TI antibodies that we describe here are encoded by diverse V_H_ genes and have differing HCDR3 lengths, and would not be detected by repertoire sequencing focused solely on heavy chain genes. Instead of the typical heavy-chain-driven interaction, the mode of molecular recognition for this class of antibodies may be determined instead mostly by canonical features of the light chain interaction, which is less commonly examined in immune repertoire sequencing efforts. Light chains do however modulate some antibody interactions with microbial pathogens. For instance, the VRC01 class of HIV-specific antibodies often have a motif that includes a 5-residue LCDR3 and a short and flexible LCDR1^39–42^. Importantly, recent work has identified a public clonotype targeting the Ebola virus glycoprotein (GP) that is elicited after natural infection or vaccination^43,44^. These GP-specific antibodies are encoded by *IGHV3-15* and *IGLV1-40*, make conserved light chain and heavy chain contacts with GP, and have been isolated from multiple individuals. Remarkably, these Ebola-specific mAbs also exhibit convergent somatic mutations in the LCDR2 across multiple donors, suggesting that the TI epitope recognition motif we describe here may be one example of a broader class of public clonotypes in which both heavy chain and light chain motifs play important roles in epitope recognition. As paired sequencing of heavy and light chain variable genes in immune repertoires becomes increasing common, it is likely that similar public clonotypes that target epitopes from a variety of pathogens will be identified.

In summary, we identified the genetic and structural basis for recognition of the influenza virus HA head domain trimer interface by human antibodies that are easily elicited in diverse individuals. The sequence studies reveal a canonical motif comprising residues in the heavy and light chain from which we can infer TI-specificity. This TI class of antibodies exhibits broad heterosubtypic binding, and lineages of TI antibodies can acquire even broader or near universal recognition of influenza type A viruses. The antibodies disrupt HA trimers, and they protect against influenza replication and disease *in vivo*. The breadth and protective capacity of the antibodies are remarkable, since very few somatic mutations are required to achieve broad recognition of influenza A strains. Furthermore, the common appearance of this class of antibody in diverse individuals suggests that the functional role of antibodies targeting the TI antigenic site in protection from seasonal influenza should be explored further.

## DATA AVAILABILITY

Atomic coordinates and structure factors for the crystal structures of H5.28 and H5.31 Fabs in their complexes with HA head domains have been deposited in the Protein Data Bank with the accession codes 6P3S (rH5.28 Fab in complex with H5 head domain), and 6P3R (rH5.31 Fab in complex with H5 head domain). The sequences of H5.28, H5.31, FluA-151, and sibling mAbs are available from GenBank.

## OBTAINING BIOLOGICAL MATERIALS

Further information and requests for reagents may be directed to and be fulfilled by the corresponding author: James E. Crowe, Jr. Materials described in this paper are available for distribution under the Uniform Biological Material Transfer Agreement, a master agreement that was developed by the NIH to simplify transfers of biological research materials.

## SUPPLEMENTARY INFORMATION

Supplementary information including supplementary experimental procedures, figures, and tables can be found with this article online.

## AUTHOR CONTRIBUTIONS

S.J.Z., J.D., I.G., N.T., and J.E.C., conceived and designed the research; S.J., I.G., P.G., N.J.T., N.K., C.S. and J.E.C. isolated, sequenced and analyzed H5.28,d H5.31, FluA-151 and the clonally related antibodies; S.J.Z., I.G., P.G., N.J.T., S.B., and S.L. performed *in vitro* profiling of mAb activities; I.G., P.G., and J.E.C. designed and analyzed mouse studies on the *in vivo* activity of mAbs; P.G., R.P.I., N.C.S. and J.B.W. performed mouse studies; J.D. determined the X-ray structures of the Fabs and related complexes; S.L. performed the HDX-MS experiments; J.A.F., R.B., C.S. performed and interpreted sequencing experiments; R.N and R.H.C. prepared recombinant antibodies and viral proteins. H.L.T and A.B.W. performed EM studies. S.J.Z. and J.E.C. wrote the manuscript.

## ACKNOWLEDGEMENTS

This work was supported by grants from the National Institutes of Health (NIH) U19 AI117905, and NIH contract HHSN272201400024C. The project described was supported by CTSA award No. UL1 TR002243 from the National Center for Advancing Translational Sciences (NCATS). S.J.Z was supported by NIH training grant T32 AI095202. Vanderbilt University Medical Center has used the non-clinical and pre-clinical services program offered by the Division of Microbiology and Infectious Diseases (DMID) in the National Institute of Allergy and Infectious Diseases (NIAID). Funding for the H5.28 and H5.31 mouse studies research was provided by contract number HHSN272201700041I, Task Order A09, from the Respiratory Diseases Branch, DMID, NIAID, NIH, USA. The contents of this publication are solely the responsibility of the authors and do not necessarily represent the official views of NCATS, NIAID or NIH. X-ray diffraction data were collected at Beamline 21-ID-G at the Advanced Photon Source, a U.S. Department of Energy (DOE) Office of Science User Facility operated for the Office of Science by Argonne National Laboratory under Contract No. DE-AC02-06CH11357. Use of the LS-CAT Sector 21 was supported by the Michigan Economic Development Corporation and the Michigan Technology Tri-Corridor (Grant 085P1000817). Support for crystallography was provided from the Vanderbilt Center for Structural Biology.

---

**Extended Data Fig. 1.**
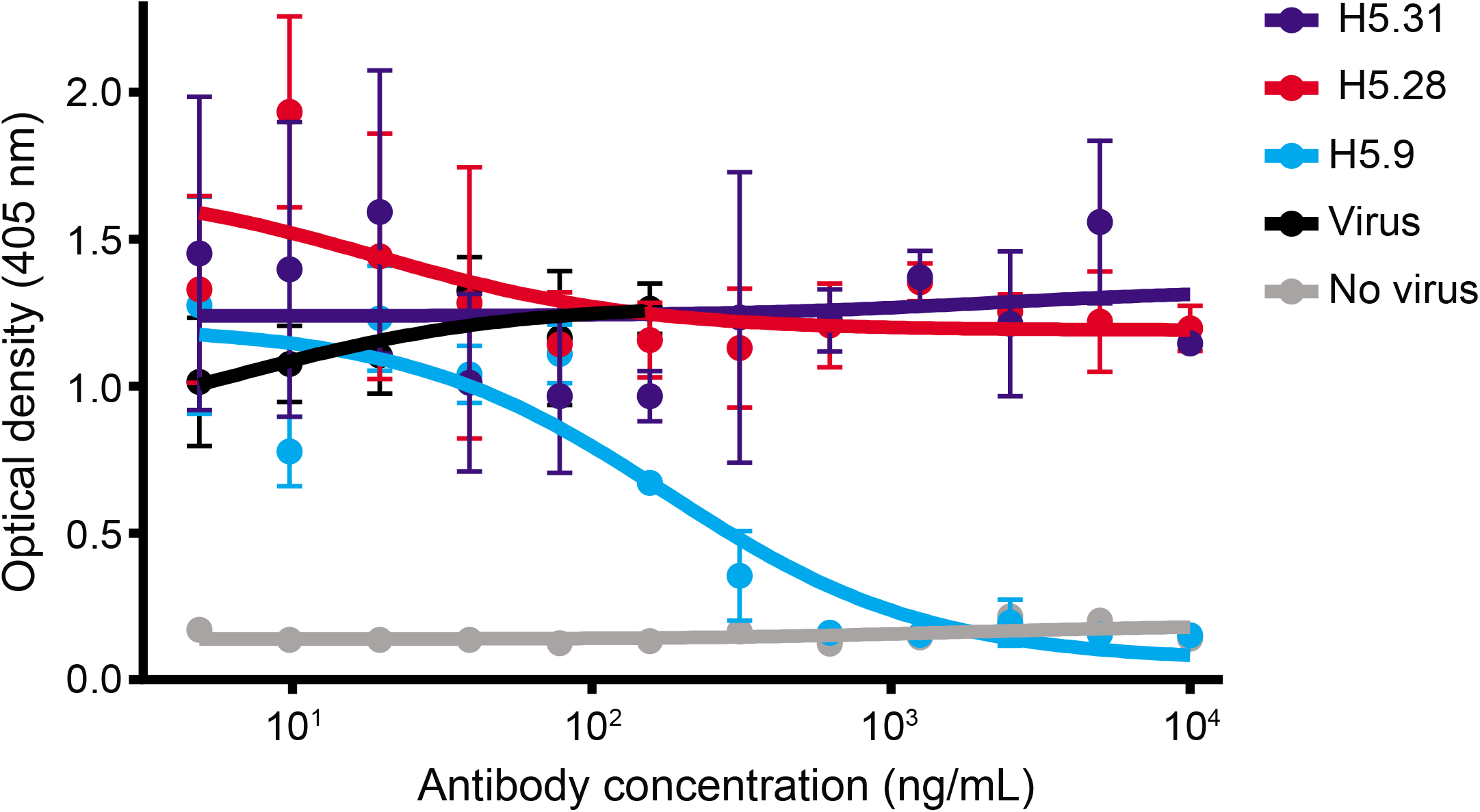
H5.31 and H5.28 do not neutralize influenza virus. The neutralization activities of H5.31 and H5.28 mAbs were tested against rgA/Vietnam/1203/2004 by a microneutralization assay on MDCK cells. Human mAb H5.9 used as positive control. Data represent the mean and SD values of technical triplicates.

**Extended Data Fig. 2.**
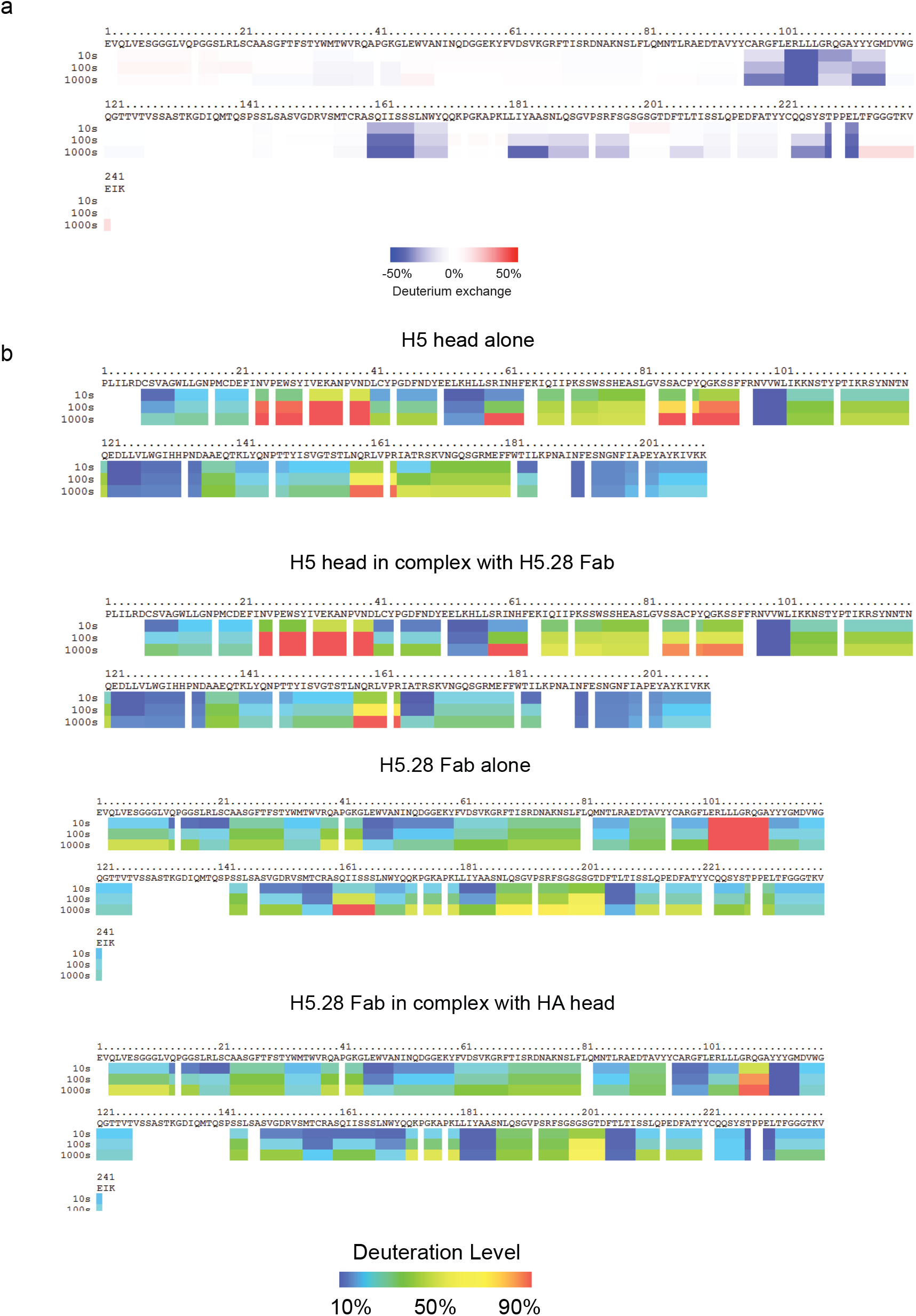
HDX data for H5.28 Fab and H5 HA monomer. **a**, The amino acid sequence of the H5.28 Fab is shown with a ribbon diagram indicating differences in deuterium uptake between the Fab alone and the Fab complexed with H5 HA monomeric head domain. Blue colors indicating slower deuterium exchange in the presence of binding to H5 HA, while red colors indicate faster deuterium exchange in the presence of H5 HA. Data are shown for 10s, 100s, and 1000s of deuterium labeling. **b**. Ribbon diagrams of deuteration level for H5.28 alone, H5 HA monomeric head domain alone, and the H5.28:H5 HA head complex. HA residues are numbered from the start of the monomeric H5 HA head construct.

**Extended Data Fig. 3.**
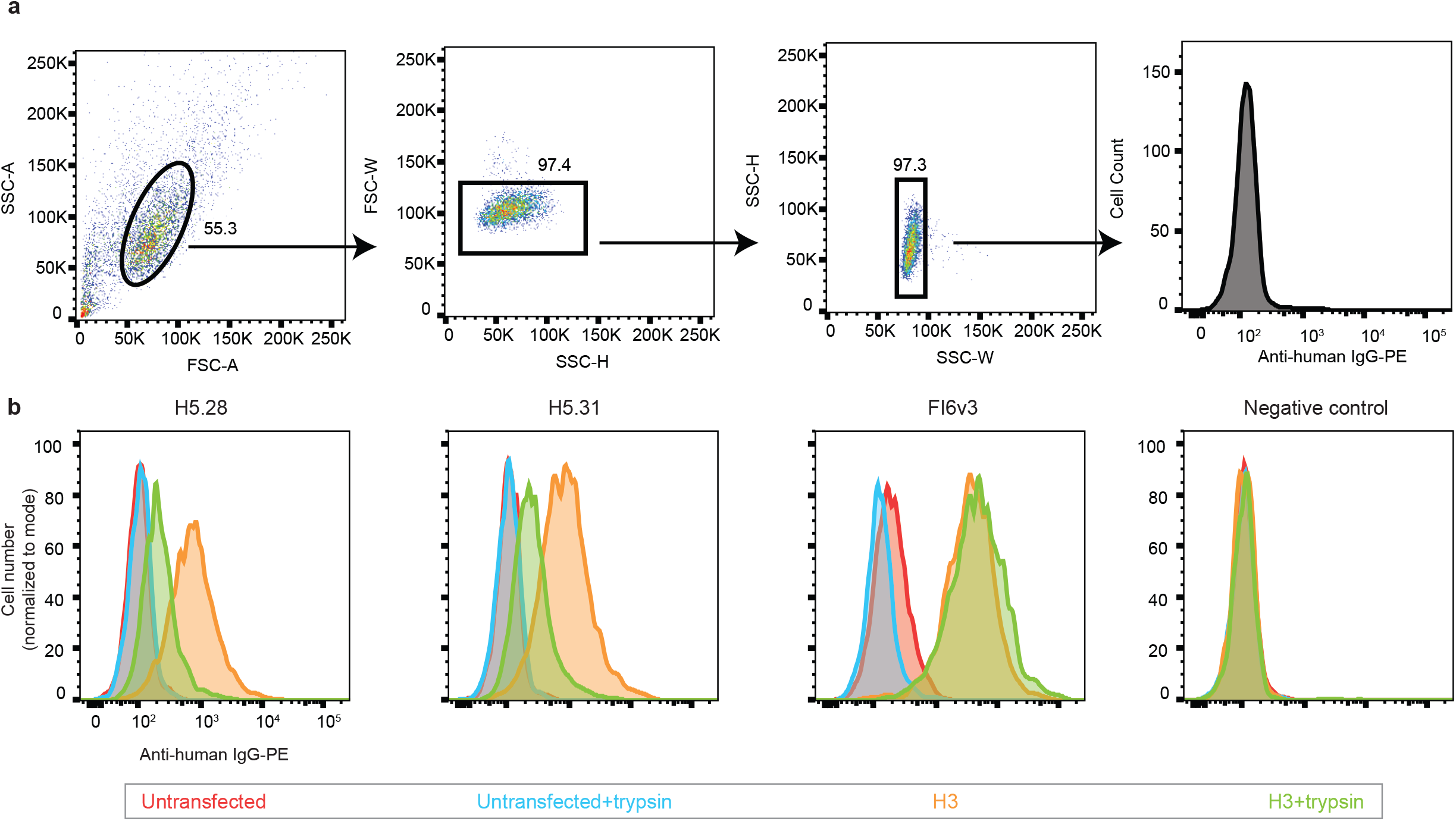
Flow cytometry gating for TI mAb binding assay using cell surface-displayed HA. **a,** Representative gating strategy shown for cells stained with a negative-control antibody. Arrows indicate which population each plot is derived from, while numbers indicate percentages of cells within each gate. **b**, Histograms showing the binding of H5.28, H5.31, and FI6v3 to untransfected cells (red), untransfected cells treated with trypsin (blue), H3 HA-transfected cells (green), and H3 HA-transfected cells treated with trypsin (orange). For each sample up to 6,000 events were collected. Data shown are a representative replicate of experimental triplicates.

**Extended Data Fig. 4.**
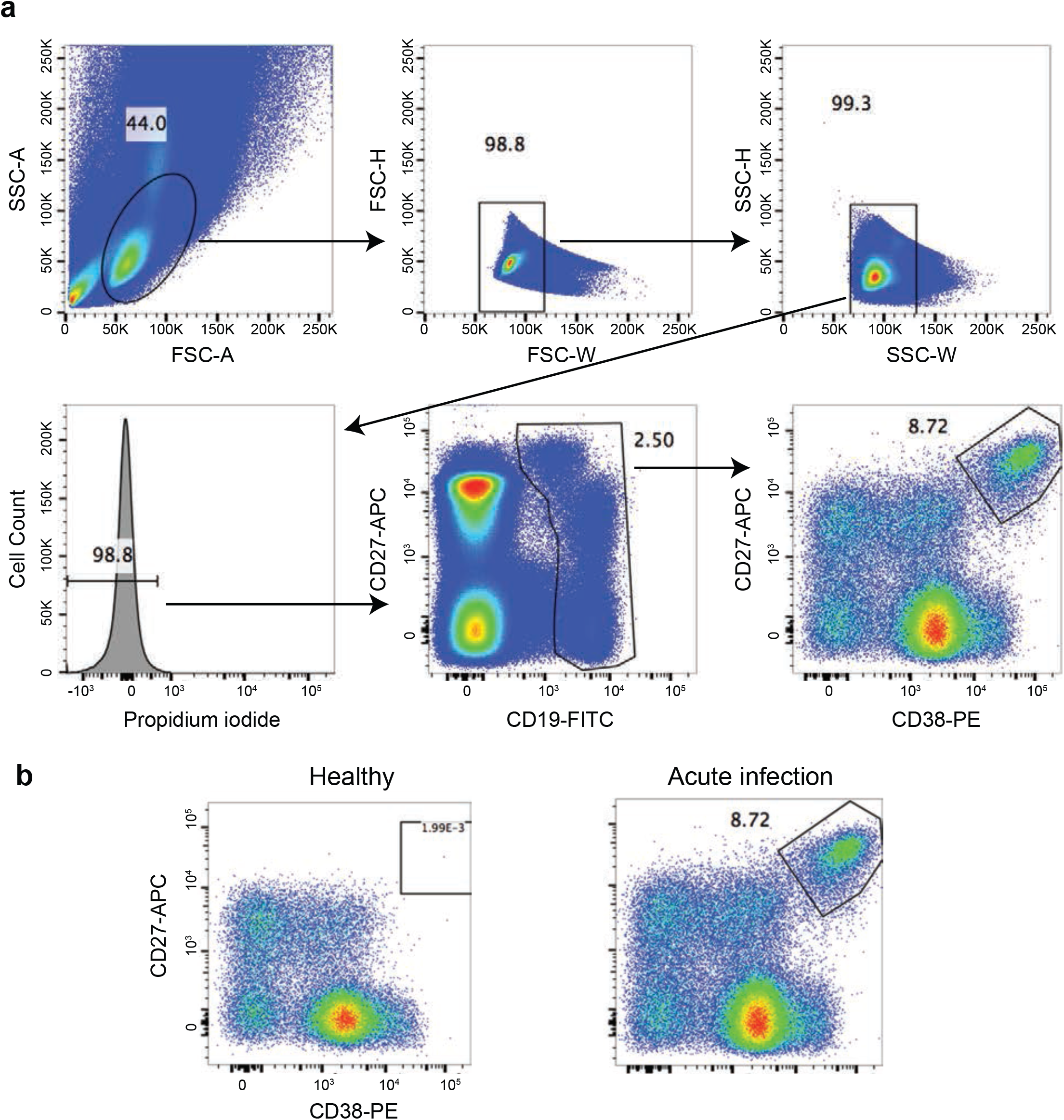
Sorting of plasmablasts from PBMCs of an influenza-infected donor. **a**, The gating strategy for isolation of plasmablasts from purified PBMCs is shown. Arrows indicate the gate that each plot is derived from, while numbers indicate the percentage of cells within each gate relative to the total number of cells. Approximately 2.2×10^7^ PBMCs were stained with CD19-FITC, CD27-APC, and CD38-PE antibodies, After gating for singlets and viable cells, plasmablasts were identified as a CD19^low^ CD27^high^ CD38^high^ population and sorted for sequencing. **b**, The CD19^low^ CD27^high^ CD38^high^ plasmablast population is absent in a healthy timepoint from the same donor (left), but is abundant 7 days after symptom onset (right).

**Extended Data Fig. 5.**
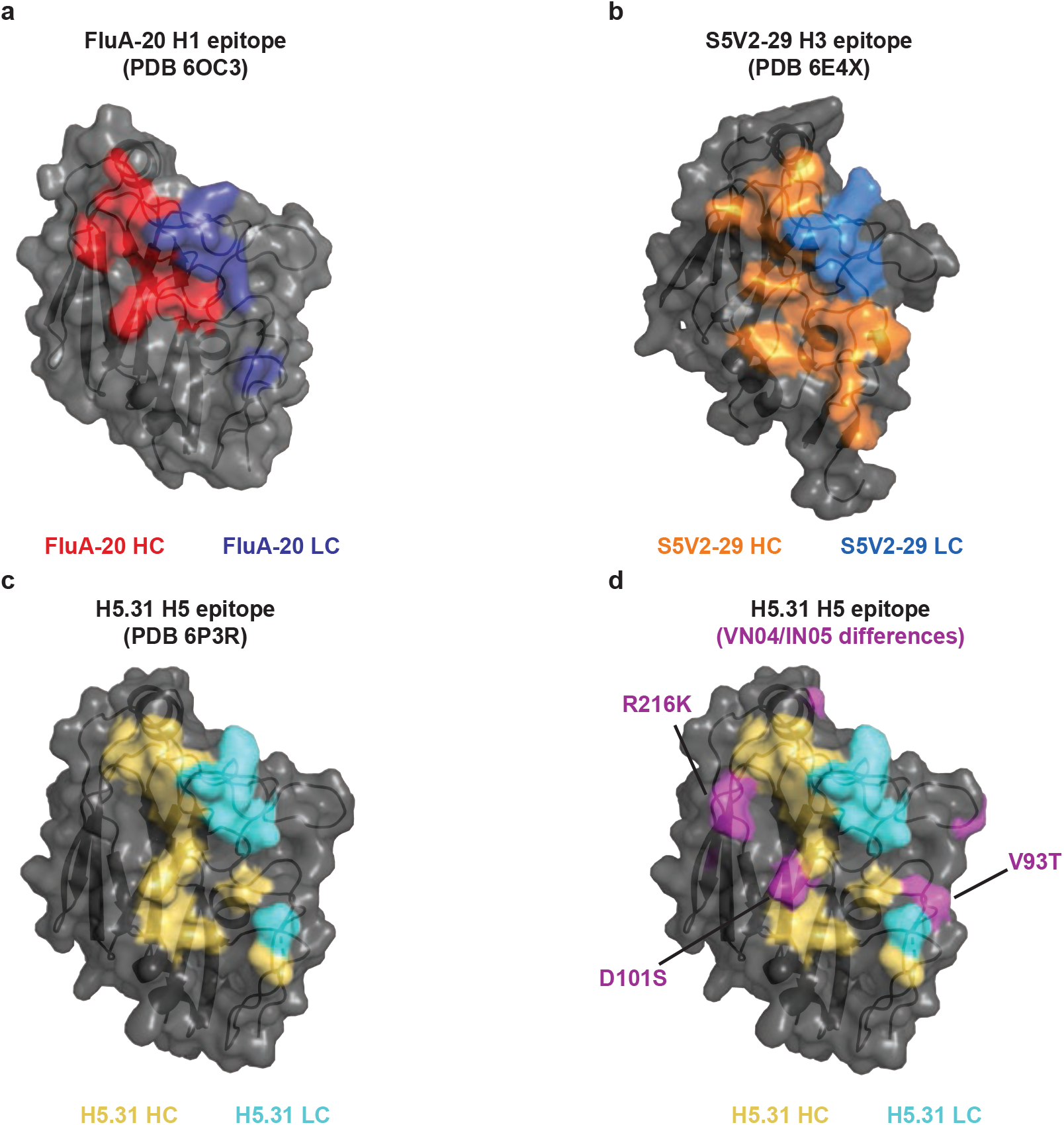
Analysis of contact residues of FluA-20, S5V2-29, and H5.31. HA residues within 3.9 Å of antibody heavy or light chains are shown for (**a**) FluA-20 in complex with H1 HA (6OC3), (**b**) S5V2-29 in complex with H3 HA (6E4X) and (**c**) H5.31 in complex with H5 HA (6P3R). Residues are colored based on their proximity to either the heavy chain or the light chain for each mAb. **d**, the H5.31 epitope with differences between VN/04 and IN/05 HAs shown in purple, illustrating putative residues responsible for the reduced binding H5.28 and H5.31 to IN/05 HA.

**Extended Data Fig. 6.**
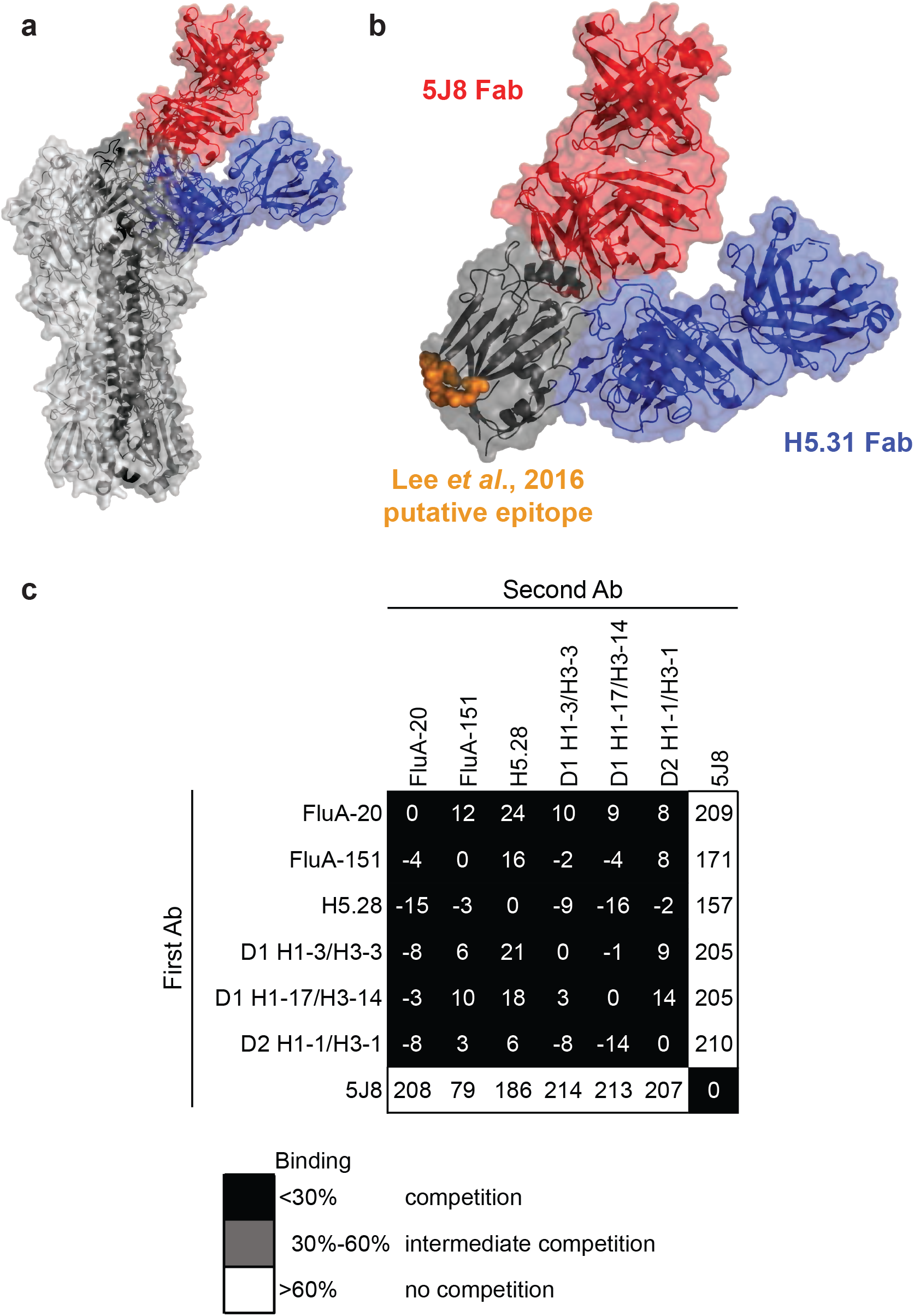
Mapping of previously reported mAbs to the same trimer-interface epitope. **a**, Structural alignment showing the H5.31:H5 HA head structure (PDB 6P3R) and the 5J8:H1 HA head structure (4M5Z) aligned with the A/California/04/2009 trimer (PDB 3UBN). The TI-specific H5.31 Fab is colored blue, while the receptor binding site (RBS) specific 5J8 Fab is colored red, and the individual protomers of the HA trimer are colored in shades of grey. **b**, Structural alignment of H5.31 and 5J8 structures with monomeric H1 head domain. Residues colored in orange illustrate the previously suggested epitope of mAbs D1 H1-3/H3-3, D1 H1-17/H3-14, and D2 H1-1/H3-1 (from Lee *et al*, 2016). **c**, biolayer interferometry-based competition assay with A/California/07/2009 H1 monomeric head domain as the antigen. TI mAbs strongly competed with one another and with D1 H1-3/H3-3, D1 H1-17/H3-14, or D2 H1-1/H3-1 for binding to H1 monomeric head domain, but did not compete with mAb 5J8, which binds the HA RBS.

## METHODS

### Expression of soluble HA proteins for binding studies

Sequences encoding the HA genes of interest were optimized for mammalian cell expression, and cDNAs were synthesized (Genscript) as soluble trimeric constructs as described previously ^45^. A monomeric HA head domain construct was synthesized with an HA-derived signal peptide sequence, an N-terminal 6-His tag, an AviTag site-specific biotinylation sequence, a thrombin cleavage site, and residues 52-263 of the A/California/07/2009 H1 HA head domain. HA proteins were expressed by transient transfection of 293F (ThermoFisher Scientific, R79007) or Expi293F (ThermoFisher Scientific, A1452) cells and grown in expression medium (ThermoFisher Scientific). Cell supernatants were harvested after 7 days, sterilized by filtration with a 0.4 μm filter and recombinant proteins were purified with HisTrap TALON FF crude or HisTrap Excel columns (GE Healthcare Life Sciences).

### PBMC and plasmablast isolation and repertoire sequencing

Studies were approved by the Vanderbilt University Medical Center Institutional Review Board. Peripheral blood was collected from a healthy donor with prior history of many seasonal influenza vaccinations, H5N1 vaccination, and H7N9 vaccination after written informed consent. For longitudinal repertoire sequencing, PBMCs from the donor were isolated by density gradient separation on Ficoll, cryopreserved and stored in liquid nitrogen storage until use. Total RNA was extracted from 10 million PBMCs. In some instances, a one-step RT-PCR was performed for 25 cycles using heavy chain BIOMED-2 variable antibody gene-specific primers as previously described ^45–47^ and the OneStep SuperScript III with Platinum® Taq High Fidelity kit (Invitrogen, 11304011).

The Illumina-specific adapters were added using the Illumina TruSeq Library Preparation Kit (Illumina, FC-121-3001) according to the manufacturer’s recommendations. The final amplicon libraries were sequenced on an Illumina MiSeq instrument using the MiSeq PE-300 v3 reagent kit (Illumina, MS-102-3001). Sequence analysis was performed using IG-BLAST v1.4, and results were parsed to MongoDB for further study. In other instances, we followed a previously described 5’ RACE approach incorporating unique molecular identifiers (UMIs) for bulk unpaired B cell repertoire sequencing ^48^. Final libraries generated using this approach were sequenced in a symmetric (r1:300 cycles and r2: 300 cycles) or asymmetric (r1:30 cycles and r2: 270 cycles) fashion using the MiSeq PE-300 v3 reagent kit (Illumina, MS-102-3001) or NovaSeq 6000 S1 reagent kit (Illumina, 20012863), respectively. For sequencing the plasmablast response to H3N2 infection, PBMCs were isolated upon natural H3N2 infection on day 7 from symptom onset. Approximately 2.2×10^7^ PBMCs were stained in FACS buffer (D-PBS supplemented with 2% FBS and 1mM EDTA) with the following phenotyping antibodies; anti-CD19-FITC (1:20 dilution, eBioscience, 11-0199-42), anti-CD27-APC (1:20 dilution, BD Biosciences, 558664), and anti-CD38-PE (1:25 dilution, BD Biosciences, 555460). Cells were resuspended in sc-V_H_:V_L_Seq sequencing buffer (D-PBS supplemented with 0.04% non-acetylated BSA) containing propidium iodide as a viability dye. Approximately 28,000 viable CD19^Low^ CD27^high^ CD38^high^ cells were sorted into sc-V_H_:V_L_Seq sequencing buffer. ~20,000 plasmablasts were carried through single-cell RNA sequencing using the 10X Genomics Chromium platform with enrichment using the 5’ VDJ amplification kit (10X Genomics) according to manufacturer instructions. Amplicons were sequenced on an Illumina Novaseq 6000, and data were processed using the CellRanger software v3.1.0 (10X Genomics). cDNA encoding heavy and light chains of interest were synthesized and cloned into IgG1 and IgK/IgL expression vectors, respectively (Twist Bioscience). Heavy and light chain plasmids were transfected into 96-well cultures of ExpiCHO cells (ThermoFisher Scientific, A29127) for microscale expression. After 7 days, mAbs were purified using protein G sepharose resin (GE Life Sciences) and buffer-exchanged into D-PBS using Zeba 96-well desalting plates (Thermo-Fisher Scientific).

### Generation of H5.28 and H5.31 hybridomas and purification of IgG

PBMCs were isolated from a donor who had received experimental H5N1 vaccines, as previously described ^30^. Briefly, human B cells in the PBMC suspension were immortalized by transformation with EBV in the presence of CpG10103, cyclosporin A, and a Chk2 inhibitor and plated in 384-well culture plates. On day 8, the supernatants from transformed B cells were used to screen for binding to recombinant H5 HA (A/Vietnam/1204/2005). The selected cloned cell lines secreting mAb H5.28 or H5.31 were grown initially in hybridoma growth medium (ClonaCell-HY medium E from STEMCELL Technologies). Prior to antibody expression and purification, these cell lines were switched to serum-free medium (GIBCO Hybridoma-SFM, Invitrogen). IgG from the hybridoma cell line supernatants was purified by affinity chromatography using protein G columns (GE Life Sciences). Purified H5.28 and H5.31 IgG generated from hybridomas was used for EC50 binding measurements, competition-binding assays, and animal studies.

### Identification of FluA-151 siblings and phylogenetic analysis

Using a database of curated antibody sequences from the FluA-151 donor, we searched CDR amino acid sequences in sequences encoded by both of the inferred germline genes for FluA-151 (*IGHV3-30* and *IGHJ4* for the heavy chain; *IGKV1-39* and *IGKJ1* for the light chain). From this pool of sequences, we selected heavy and light chains with a CDR3 length between 19 and 13 amino acids for the heavy chain and 10 and 8 amino acids for the light chain. We then ran blastp with these CDR3s, the CDR3s of FluA-151, and the seven FluA-151-related sequences identified in the single-cell RNA sequencing to obtain values for percent coverage and percent identity. We averaged the percent coverage and percent identity values to score these sequences and designated sequences with scores >85% against FluA-151 or any of the other seven plasmablast-derived sequences as FluA-151 siblings. For these siblings, we extracted the full-length nucleotide sequences and aligned those sequences to the corresponding germline gene (*IGHV3-30*18* or *IGKV1-39*01*) as well as FluA-151 and FluA-151 sibling sequences using Clustal Omega v1.2.0. We used the PHYLIP phylogenetic software package v3.697 to generate a maximum-likelihood tree from the aligned sequences using the DNAML program, using the sequence of the germline *IGHV* or *IGKV* gene as an outgroup. The resulting phylogenetic trees were visualized using the FigTree phylogenetic tree viewer (FigTree v1.4.4) to color branches corresponding to sequencing timepoints, and the heavy and light chain sequences of the FluA-151 inferred common ancestor (UCA) were extracted from the PHYLIP-generated tree.

### Sequencing of antibody genes from hybridomas

Antibody heavy and light chain variable region genes were sequenced from antigen-specific hybridoma lines that had been cloned biologically from flow cytometry. Total RNA was extracted using the RNeasy Mini kit (Qiagen).We modified a previously described 5’ RACE approach for target enrichment and sequencing ^48^. Briefly, 5 μL total RNA was mixed with cDNA synthesis primer mix (10 μM each) and incubated for 2 min at 70°C and then decrease the incubation temperature to 42°C to anneal the synthesis primers (1-3 min). After incubation, a mix containing 5X first-strand buffer (Clontech), DTT (20 mM), 5’ template switch oligo (10 μM), dNTP solution (10 mM each) and 10X SMARTScribe Reverse Transcriptase (Clontech) was added to the primer-annealed total RNA reaction and incubated for 60 min at 42°C. The first-strand synthesis reaction was purified using the Ampure Size Select Magnetic Bead Kit at a ratio of 0.6X (Beckman Coulter). Following, a single PCR amplification reaction containing 5 μL first-strand cDNA, 2X Q5 High Fidelity Mastermix (NEB), dNTP (10 mM each), forward universal primer (10 μM) and reverse primer mix (0.2 μM each in heavy chain mix, 0.2 μM each in light chain mix) were subjected to thermal cycling with the following conditions: initial denaturation for 1 min 30 s followed by 30 cycles of denaturation at 98°C for 10 s, annealing at 60°C for 20 s, and extension at 72°C for 40 s, followed by a final extension step at 72°C for 4 min. The first PCR reaction was purified using the AMPure Size Select Magnetic Bead Kit at a ratio of 0.6X (Beckman Coulter). Amplicon libraries were then prepared according to the Pacific Biosciences Multiplex SMRT Sequencing protocol and sequenced on a Pacific Biosciences Sequel platform (Pacific Biosciences, Menlo Park, CA). Raw sequencing data was demultiplexed and circular consensus sequences (CCS) were determined using the Pacific Biosciences SMRT Analysis tool suite v8.0. The identities of gene segments and mutations from germlines were determined by alignment using ImMunoGeneTics database ^49,50^.

### ELISAS and determination of half maximal effective concentration (EC_50_) for binding

ELISAs were performed using 384-well plates that were coated overnight at 2 μg/mL with the recombinant HA protein of interest. The plates then were blocked with 50 μL of 5% non-fat dry milk and 0.1% Tween-20 in D-PBS (ELISA buffer) for 1 hr at RT. The plates were washed and three-fold dilutions of the mAb in ELISA buffer at a starting concentration of 10 μg/mL were added to the wells and incubated for an hour. The plates were washed and 25 μL of ELISA buffer containing a 1:4,000 dilution of anti-human IgG alkaline phosphatase conjugate (Meridian Life Science, W99008A) was added. After a final wash, 25 μL of phosphatase substrate solution (1 mg/mL p-nitrophenol phosphate in 1 M Tris aminomethane) was added to the plates, incubated for 1 hr and the optical density values were measured at 405 nm wavelength on a BioTek plate reader. The plates were washed 3 times between each step with PBS containing 0.05% Tween-20. Each dilution was performed in triplicate, and the EC50 values were calculated in Prism software (GraphPad) v8.1.1 using non-linear regression analysis. Each experiment was conducted twice independently.

### *In vivo* protection study for mAbs H5.28 and H5.31

To assess protective efficacy of mAbs, female 18-20 g BALB/c mice (Charles River Laboratories, Wilmington, MA) were inoculated by the intraperitoneal (i.p.) route with a 1, 3, or 10 mg/kg dose of individual mAbs. Human anti-dengue virus mAb DENV 2D22 served as a mock control treatment at dose 10 mg/kg. Oseltamivir phosphate (hereafter referred to as oseltamivir) (Roche, Palo Alto, CA) diluted in sterile PBS was inoculated i.p. at 30 mg/kg/day and served as a positive control. In ABSL-2 facilities, ketamine-xylazine anesthetized mice were inoculated by the intranasal (i.n.) route at 24 hours after the mAb treatment with 2,200 50% cell culture infectious doses (CCID50) of a mouse-adapted influenza A/California/04/2009 (H1N1pdm) in 90 μL of sterile PBS. Oseltamivir treatments were given i.p. twice daily for 5 days, starting at 1 h post-infection. Mice were weighed and monitored daily for body weight change and signs of disease for 21 days, and those losing over 30% of initial body weight were humanely euthanized as per IACUC requirements. This study was conducted in the AAALAC-accredited laboratory animal research center of Utah State University in accordance with the approval of the institutional animal care and use committee of Utah State University.

### *In vivo* protection study for FluA-151 WT or LALA-PG Fc variant mAb

For prophylactic studies measuring protection from weight loss following sub-lethal H1N1 challenge, groups of ten BALB/c mice were given 10 mg/kg of FluA-151 WT, FluA-151 LALA-PG, positive control anti-stem mAb MEDI-8852, or negative control anti-dengue mAb 2D22 via the intraperitoneal route one day prior to inoculation. The following day, mice were challenged with a sublethal dose of 6.74 x 10^4^ FFU of a mouse-adapted A/California/04/2009 H1N1 strain. Mice were monitored for 14 days for weight change kinetics. Weight change curves between groups were compared using a two-way ANOVA with Tukey’s multiple comparisons correction. Studies were conducted in A-BSL2 facilities at Vanderbilt University Medical Center in accordance with the approval of the Vanderbilt University Medical Center Institutional Animal Care and Use Committee.

### Competition-binding assays

Biolayer interferometry on an Octet Red instrument (FortéBio) was used to perform competition-binding assays. Briefly, antigen and antibodies were diluted in D-PBS with 1% BSA and 0.05% Tween20. We first loaded either trimeric recombinant HA from H1 A/California/04/2009 or monomeric HA head domain from H1 A/California/07/2009 onto Ni-NTA tips at a concentration of 20 μg/mL. We then tested binding of two successively applied mAbs at 50 μg/mL. Competition was analyzed using the Octet analysis software (Data Analysis 9, FortéBio). Binding values were normalized to the binding signal measured in the absence of the first antibody, and self-self competition values were subtracted. Antibodies were defined as competing antibodies if the first antibody reduced binding of the second antibody by more than 70 percent. Antibodies were defined as non-competing antibodies if the first antibody reduced binding of the second antibody by less than 40 percent. Antibodies were defined as partially competing antibodies if the first antibody reduced binding of the second antibody between 40 and 70 percent.

### Recombinant protein expression and purification for crystallography

A cDNA encoding the head domain of VN/04 HA (residues 58 - 268) was codon optimized, synthesized, and cloned into pcDNA3.1 (+) downstream of the signal peptide from pHLsec vector (Genscript). To facilitate protein purification, a linker sequence (AA) and a 6-his tag were added to the C-terminus of the construct. Synthetic cDNAs encoding the heavy chains and light chains of H5.31 and H5.28 Fabs were synthesized and cloned into pcDNA3.1 (+) downstream of the CD5 signal peptide (Genscript). Expi293F cells were transfected transiently with pcDNA3.1 (+) plasmids encoding the HA head domain, and culture supernatants were harvested after 6 to 7 days. The head domain was purified from the supernatants by nickel affinity chromatography with HisTrap Excel columns (GE Healthcare Life Sciences), and H5.31 and H5.28 Fabs were purified with CaptureSelect IgG-CH1 (Thermo Fisher Scientific). To obtain complexes of Fabs and HA head domain, purified recombinant Fabs were mixed with excess HA head domain in a molar ratio of 1:2. The mixture was incubated for ~ 10 minutes, and the complexes were purified from the mixture using a HiLoad^®^ 16/600 Superdex^®^ 200 size-exclusion column (GE Healthcare Life Sciences).

### Crystallization, data collection, and structure determination

The complexes were concentrated to ~ 10 mg/mL in a buffer of 20 mM Tris-HCl pH 7.5, 50 mM NaCl, and then the concentrated samples were used for crystallization screen and optimization. Crystals of the complexes (H5.28-HA and H5.31-HA) were grown by the vapor diffusion method. Extensive initial crystallization screening was carried out with a TTP Labtech Mosquito robot (TTP Labtech), and subsequent crystallization optimization was performed manually using hanging-drop vapor diffusion method with 15-well screw-cap crystallization plates (Qiagen). The H5.28-HA head complex was crystallized in 7-9% PEG 8000, 0.3-0.7 M calcium acetate, 0.1 M imizadole pH 8.0, and the H5.31-HA head domain in 0.8 M potassium phosphate dibasic, 0.6 M sodium phosphate monobasic. The crystals were flash frozen in liquid nitrogen with 30% glycerol as the cryoprotectant. Diffraction data were collected at the Advanced Photon Source LS-CAT beamline 21-ID-G. The data were processed and integrated with XDS data processing software ^51^, and scaled with the software SCALA^52^. The crystal structures of the both complexes were solved by molecular replacement using the crystal structure of VN/04 HA head domain (PDBID: 4XNQ) and the crystal structure of human anti-Marburg Fab MR78 (PDB ID: 5JRP) as the searching models with the program Phaser^53^. The structures were refined and manually rebuilt with Phenix^54^ and Coot^55^. The data collection and refinement statistics are shown in Supplementary Table 1. Structure figures were generated by MacPyMOL (DeLano Scientific LLC).

### Peptide fragmentation and deuterium exchange mass spectrometry

To maximize peptide probe coverage, the optimized quench condition was determined prior to deuteration studies^56,57^. In short, the HA head domain was diluted with buffer of 8.3 mM Tris, 150 mM NaCl, in H_2_O, pH 7.15) at 0 °C and then quenched with 0.8% formic acid (v/v) containing various concentration of GuHCl (0.8 - 6.4 M) and Tris(2-carboxyethyl)phosphine (TCEP) (0.1 or 1.0 M). After incubating on ice for 5 min, the quenched samples were diluted 4-fold with 0.8% formic acid (v/v) containing 16.6% (v/v) glycerol and then were frozen at −80°C until they were transferred to the cryogenic autosampler. Using the quench buffer of 1.4 M GuHCl, 100 mM TCEP in 0.8% formic acid gave an optimal peptide coverage map. The samples later were thawed automatically on ice and then immediately passed over an AL-20-pepsin column (16 μL bed volume, 30 mg/mL porcine pepsin (Sigma)). The resulting peptides were collected on a C18 trap and separated using a C18 reversed phase column (Vydac) running a linear gradient of 0.046% (v/v) trifluoroacetic acid, 6.4% (v/v) acetonitrile to 0.03% (v/v) trifluoroacetic acid, 38.4% (v/v) acetonitrile over 30 min with column effluent directed into an Orbitrap Elite mass spectrometer (Thermo-Fisher Scientific). Data were acquired in both data-dependent MS:MS mode and MS1 profile mode. Proteome Discoverer software (Thermo Finnigan Inc.) was used to identify the sequence of the peptide ions. DXMS Explorer (Sierra Analytics Inc., Modesto, CA) was used for the analysis of the mass spectra as described previously ^58^. Fab bound HAs were prepared by mixing H5.28 Fab with monomeric H5 head domain at a 1:1.3 (HA:H5.28 Fab) or 1.2:1 (HA:H5.28 Fab) stoichiometric ratio. The mixtures were incubated at 25 °C for 30 min. All functionally deuterated samples (with the exception of the equilibrium-deuterated control) and buffers were pre-chilled on ice and prepared in the cold room. Functional deuterium-hydrogen exchange reaction was initiated by diluting free HA or antibody-bound HA stock solution with D_2_O buffer (8.3 mM Tris, 150 mM NaCl, in D_2_O, pDREAD 7.15) at a 1:2 vol/vol ratio. At 10 sec, 100 sec and 1,000 sec, the quench solution was added to the respective samples, and then incubated on for 5 minutes before frozen at −80 °C. In addition, nondeuterated samples, equilibrium-deuterated back-exchange control samples were prepared as previously described 56,5^7^,59. The centroids of the isotopic envelopes of nondeuterated, functionally deuterated, and fully deuterated peptides were measured using DXMS Explorer, and then converted to corresponding deuteration levels with corrections for back-exchange^60^.

### Influenza viruses

The virus stocks used for neutralization assays were propagated in MDCK cells (American Type Culture Collection, CCL-34). Supernatants were collected from virus-infected MDCK cell culture monolayers in plain Dulbecco’s Modified Eagle Medium supplemented with 2 μg/mL of TPCK-trypsin. The virus used for murine experiments, a mouse-adapted A/California/04/2009 H1N1 strain, was propagated in embryonated chicken eggs.

### Microneutralization assays

Neutralization potential of H5.28 and H5.31 were determined by microneutralization and HAI assays, as previously described^45^. Briefly, 2-fold serial dilutions of each antibody in viral growth medium (plain DMEM supplemented with 2 μg/mL of TPCK-trypsin and 50 μg/mL gentamicin). were mixed with an equivalent volume of viral growth medium containing 100 TCID50 of virus. Antibodies and virus were incubated for 1 hr at RT The MDCK cell monolayer cultures were washed twice with 100 μL PBS containing 0.1% Tween-20, and the virus-antibody mixture then was added to cells and incubated for 32 hours at 37°C. After incubation, cells were washed and fixed with 100 μL of 80% methanol/20% PBS. The presence of influenza nucleoprotein in the fixed cells was determined by ELISA using a 1:8,000 dilution of mouse anti-NP antibody (BEI Resources, NR-4282) as the primary antibody and a 1:4,000 dilution of goat anti-mouse alkaline phosphate conjugate as the secondary antibody (ThermoFisher Scientific). Each dilution was tested in triplicate and neutralization curves were graphed using GraphPad Prism.

### Flow cytometric analysis of antibody binding to cell-surface expressed HA

HEK293F cells grown in expression medium were transfected transiently with cDNA encoding H3 A/Hong Kong/1/1968 HA protein and incubated at 37°C for 36 h. Untransfected (UT) or transfected cells were washed and incubated with either DMEM containing TPCK trypsin (2 μg/mL) or plain DMEM for 15 min at 37°C. After incubation cells were washed with PBS containing 2% of heat-inactivated FBS and 2 mM EDTA (FACS buffer). Cells then were stained with mAbs H5.28, H5.31, or FI6v3 (10 μg/mL) for 30 min at RT and for 5 min at 37°C. The cells were washed with FACS buffer and incubated with secondary goat anti-human IgG PE antibody (Southern Biotech) for 1 hour at 4°C, fixed with 4% formaldehyde in PBS, and analyzed by flow cytometry using an LSR-2 cytometer (BD Biosciences). Data for a total of up to 6,000 cell events were acquired and flow cytometry data were analyzed with FlowJo software v10.4 (Tree Star).

### Negative stain electron microscopy

H5.38 or H5.31 Fabs were incubated with uncleaved H1 HA trimer for 20 sec at 5X molar excess of Fab. The complex was added to carbon-coated 400 mesh cooper grids and stained with 2% uranyl formate. Micrographs were collected on a 120kv Tecnai Spirit microscope with a 4kx4k TemCam F416 camera using Leginon^61^. Images then were processed with Appion ^62^. Particles were selected with DoGpicker^63^, and 2D classes were generated with MSA/MRA ^64^. Particles were false colored in Photoshop CS6 (Adobe).

---

**Supplementary Table 1.** Data collection and refinement statistics for the crystals of H5.31/H5-HA and H5.28/H5-HA complexes

| **Data collection** |  |  |
| --- | --- | --- |
| Crystal | H5.31/H5-HA | H5.28/H5-HA |
| PDB ID | 6P3R | 6P3S |
| Wave length (Å) | 0.97856 | 0.97856 |
| Space group | F222 | P22<sub>1</sub>2<sub>1</sub> |
| Unit cell dimensions |  |  |
| a, b, c (Å) | 185.1, 234.5, 284.8 | 135.5, 187.2, 342.4 |
| α, β, γ | 90, 90, 90 | 90.0, 90.0, 90.0 |
| Resolution (Å) | 49.52 – 3.00 | 49.46 – 4.00 |
| Unique reflections | 61716 (8933) | 74362 (10717) |
| Redundancy | 5.9 (6.0) | 7.6 (7.7) |
| Completeness (%) | 100.0 (100.0) | 99.9 (100.0) |
| R<sub>merge</sub> (%) | 7.1 (50.8) | 21.0 (62.5) |
| I/σ(I) | 16.0 (3.5) | 8.2 (3.4) |
| **Refinement statistics** |  |  |
| R<sub>factor</sub> (%) | 19.2 | 28.1 |
| R<sub>free</sub> (%) | 22.7 | 32.4 |
| R.m.s.d. (bond) (Å) | 0.0024 | 0.0016 |
| R.m.s.d. (angle) (deg) | 0.721 | 0.545 |
| Ramachandran plot |  |  |
| Favored (%) | 95.29 | 91.95 |
| Allowed (%) | 4.47 | 7.11 |
| Outliers (%) | 0.24 | 0.94 |

$R_{\text{merge}} = \sum \sum |I_{\text{hkl}} - I_{\text{hkl}(j)}| / \sum I_{\text{hkl}}$ , where $I_{\text{hkl}(j)}$ is the observed intensity and $I_{\text{hkl}}$ is the final average intensity.
$R_{\text{work}} = \sum ||\text{Fobs}| - |\text{Fcalc}|| / \sum |\text{Fobs}|$ and $R_{\text{free}} = \sum ||\text{Fobs}| - |\text{Fcalc}|| / \sum |\text{Fobs}|$ , where $R_{\text{free}}$ and $R_{\text{work}}$ are calculated using a randomly selected test set of 5% of the data and all reflections excluding the 5% test set, respectively. Numbers in parentheses are for the highest resolution shell.

